# A Yeast Global Genetic Screen Reveals that Metformin Induces an Iron Deficiency-Like State

**DOI:** 10.1101/190389

**Authors:** B. Stynen, D. Abd-Rabbo, J. Kowarzyk, L. Miller-Fleming, M. Ralser, S.W. Michnick

## Abstract

We report here a simple and global strategy to map out gene functions and target pathways of drugs, toxins or other small molecules based on “homomer dynamics” Protein-fragment Complementation Assays (*hd*PCA). *hd*PCA measures changes in self-association (homomerization) of over 3,500 yeast proteins in yeast grown under different conditions. *hd*PCA complements genetic interaction measurements while eliminating confounding effects of gene ablation. We demonstrate that *hd*PCA accurately predicts the effects of two longevity and health-span-affecting drugs, immunosuppressant rapamycin and type II diabetes drug metformin, on cellular pathways. We also discovered an unsuspected global cellular response to metformin that resembles iron deficiency. This discovery opens a new avenue to investigate molecular mechanisms for the prevention or treatments of diabetes, cancers and other chronic diseases of aging.

## 1. Introduction

Biguanides form a class of drugs that lower blood sugar levels. Its most prominent member metformin is currently one of the most prescribed forms of non-insulin therapy among type 2 diabetes (T2D) patients. Recent investigations have revealed potential novel health benefits of metformin, in spite of the fact that its mechanism of action is still unknown. There is a considerable interest in metformin as a drug that potentially improves human health span. An observational study for humans and promising initial reports for mice on the effect of metformin on life span have led to the launch of a pioneering clinical study on the longevity effects of metformin on humans (*1-4*).

No individual specific metformin-binding protein has been identified to date and it is likely to act on multiple targets (*5, 6*). In the absence of a single target, bridging the gap between phenotype of a chemical such as metformin and identifying the cellular processes that may underlie the phenotype requires screening strategies that provide reporters for all or most known cellular processes. Ideally, this screening strategy provides information on both passive and indirect (e.g. protein/mRNA turnover) and active (e.g. post-translational modification) effects on a pathway reporter protein. High-throughput methods such as mRNA and protein abundance profiling can be informative of molecular actions of pathways. However, they measure effects of molecules indirectly and on passive processes affecting the reporters. Large-scale screening of basic eukaryotic biochemical pathways can be achieved in the simplest eukaryotic model, the budding yeast *Saccharomyces cerevisiae*, where synthetic interactions of chemicals and gene knockouts across practically all genes can be used to determine if a chemical acts in a pathway and in some cases, on specific target proteins (*7*, 8). Recently, a combination of synthetic genetic array technology with high-content screening has generated the first flux network for the budding yeast in which protein abundance and localization are tracked in response to a chemical over time (*9*).

We have developed a simple alternative to chemical epistasis in yeast that does not require gene knockout arrays and can capture integrated effects of a cellular perturbation on many reporter proteins. In our strategy, we measure condition-dependent changes in homooligomerization of proteins in living yeast using a protein-fragment complementation assay (homomer dynamics or *hd*PCA). This method is based on protein-protein interaction-driven folding and reconstitution of murine dihydrofolate reductase (mDHFR) from complementary fragments. The coding sequences of N‐ and C-terminal complementary fragments (F[1,2] and F[3]) of mDHFR are integrated into the genome 3’ to the open reading frames of genes of interest (Fig. 1A). For measurements of *hdPCA*, mDHFR F[1,2] and F[3] are integrated into the alleles of a single gene in mating type strains BY4741 *MAT***a** and BY4742 *MAT*α, respectively. Mating of these two strains results in two copies of the same gene tagged with complementary mDHFR fragments. Homo-oligomerization of the tagged protein under study brings the two mDHFR fragments together in space resulting in mDHFR folding and reconstitution of its activity. The reconstituted mDHFR is a mutant version that does not bind the cytostatic DHFR inhibitor methotrexate at concentrations that inhibit the yeast DHFR. Consequently, cells with the reconstituted mDHFR can divide in the presence of methotrexate, resulting in colonies whose size is proportional to the number of homomeric complexes per cell (*10*).

**Fig. 1.**
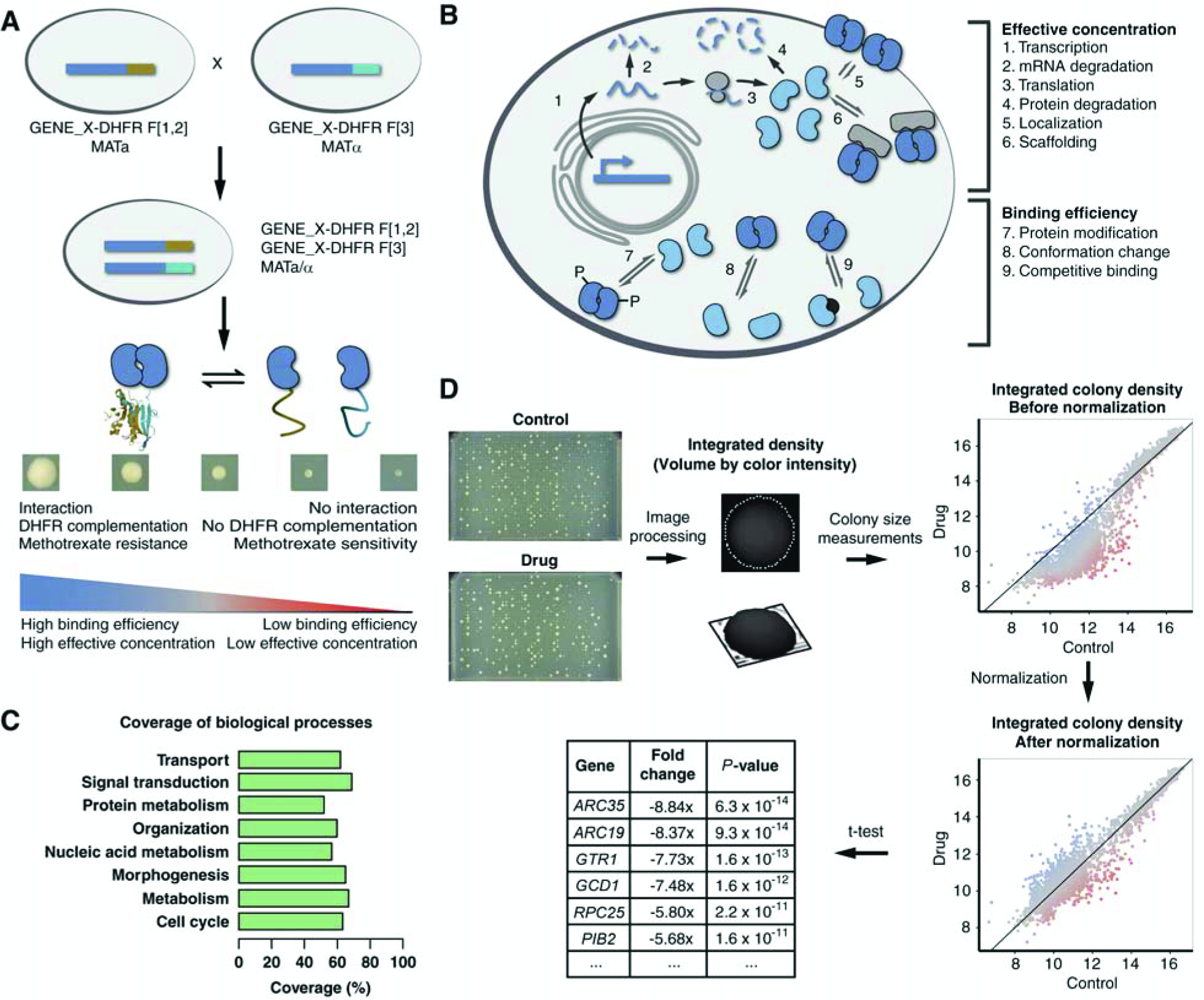
A homomer dynamics DHFR PCA (hdPCA) for the detection of the condition-dependent states of proteins. **(A)** A library of *hdPCA* strains is created by mating of two strains, each containing an ORF of interest tagged with one of the two complementary fragments of murine dihydrofolate reductase (mDHFR; brown and light blue). Upon interaction of two molecules of the same protein, the two fragments of mDHFR fold and reconstitute into a functional enzyme. This reconstitution quantitatively correlates with growth in the presence of methotrexate (*17*) and is determined by effective concentration and binding efficiency. **(B)** The degree of homomerization (self-association) of a protein is a result of different factors, some of which influence the effective concentration (upper part) while others influence binding efficiency (lower part). **(C)** Coverage of GO biological processes in the *hd*PCA, with coverage determined by the percentage of proteins associated with a GO Super-Slim biological process, that have been screened in the *hd*PCA. GO Super-Slim biological processes were obtained by manually condensing the standard terms in the GO Slim (available at http://www.yeastgenome.org/download-data/curation, as of July 2017) into eight GO global terms. **(D)** Workflow of the hdPCA.

Applications of PCA reporter screens have been reported to measure effects of drugs on specific pathways or to map out specific mechanisms of action of drugs (*11-14*). For instance, we have previously used a series of heteromeric fluorescent protein reporter PCAs to identify potential anticancer agents in mammalian cells (*12*). The hdPCA differs from these strategies because the focus is shifted from protein-protein interactions to individual protein states. Any change in the number of homomeric complexes results from the integrated passive and/or active effects of a molecule acting on a pathway in which a reporter protein participates (Fig. 1B). The change in the number of homomeric complexes will then affect the colony growth of the corresponding hdPCA strain. Examples of protein properties that the mDHFR PCA can report on include a change in binding affinity, post-translational modifications, localization and abundance (*15-18*).

Homomeric interactions are quite common; for instance 50 % of proteins in the bacterium *Mycoplasma pneumoniae* form homomers under one condition and 47 % (41,618 / 88,739) of high-resolution structures of protein complexes are homomeric (*19, 20*). It is important to note that strong binding affinities are not essential to detect changes in homomerization in the mDHFR PCA screen. This screen can report on abundance and localization of a protein even if the binding affinity between this protein and a second protein in the mDHFR PCA test is low (*17*). Likewise, changes in *hd*PCA signal of proteins with low self-binding affinity or low abundance, as indicated by their corresponding *hd*PCA colony sizes, can still be detected with statistical significance (Files S1-4). The 3,504 *hd*PCA reporters cover ~60 % of the yeast proteome equally distributed among the major categories of cellular processes (Fig. 1C). Thus, the *hdPCA* has the potential to evaluate the effects of a molecule across practically all cellular processes. In addition, the virtue of the *hdPCA* screening strategy is its simplicity and economy. The 3,504 diploid *hd*PCA strains can be arrayed at high density on just three agar plates (1,536 strains per plate) and an entire screen to test the effects of a molecule on all of the strain reporters with an appropriate number of replicates can be performed with only a dozen plates (Fig. 1D; Supplementary Methods).

*hd*PCA strains are spotted as four replicates on plates containing control or test media (*e.g*. containing drugs or toxins or missing nutrients). Next, pictures of the plates are taken on nine days within two weeks. Integrated colony density is derived from image analysis and the final values are obtained after two rounds of normalization, one to correct for plate-to-plate variability (quantile normalization) and the other to correct for the basal effect of the drug on growth (LOESS normalization). Finally, a statistically significant growth difference is determined with a regularized *t*-test that compares means of the replicates under the control and drug conditions (*21*).

## 2. hdPCA measures passive and active protein changes

To assess the performance of the *hd*PCA screen, we chose the immunosuppressant drug rapamycin as a test perturbation. We chose this molecule because (1) it has well-defined and distinct target receptors (the target-of-rapamycin (TOR) complex) and known downstream affected cellular processes, (2) like metformin, it has potential anticancer and anti-aging properties and (3) there exist a number of sources of data for large-scale effects of rapamycin on mRNA and protein homeostasis, active processes and chemical-genetic interactions (*22*). These allowed us to test the hypothesis that *hd*PCA can capture the effects of a molecule on a reporter protein and by extension on a given cellular process in which the reporter protein is implicated.

We performed the *hdPCA* screen with strain arrays grown in the presence or absence of 2 nM rapamycin. As an example of a response to the addition of rapamycin, the strain expressing the *hd*PCA for the protein Asc1 (*ASC1-DHFR-F[1,2]* and *ASC1-DHFR-F[3]*) grows significantly better in the absence of rapamycin (Fig. 2A). This reflects a change in the state of Asc1, an ortholog of human G-protein subunit RACK1.

**Fig. 2.**
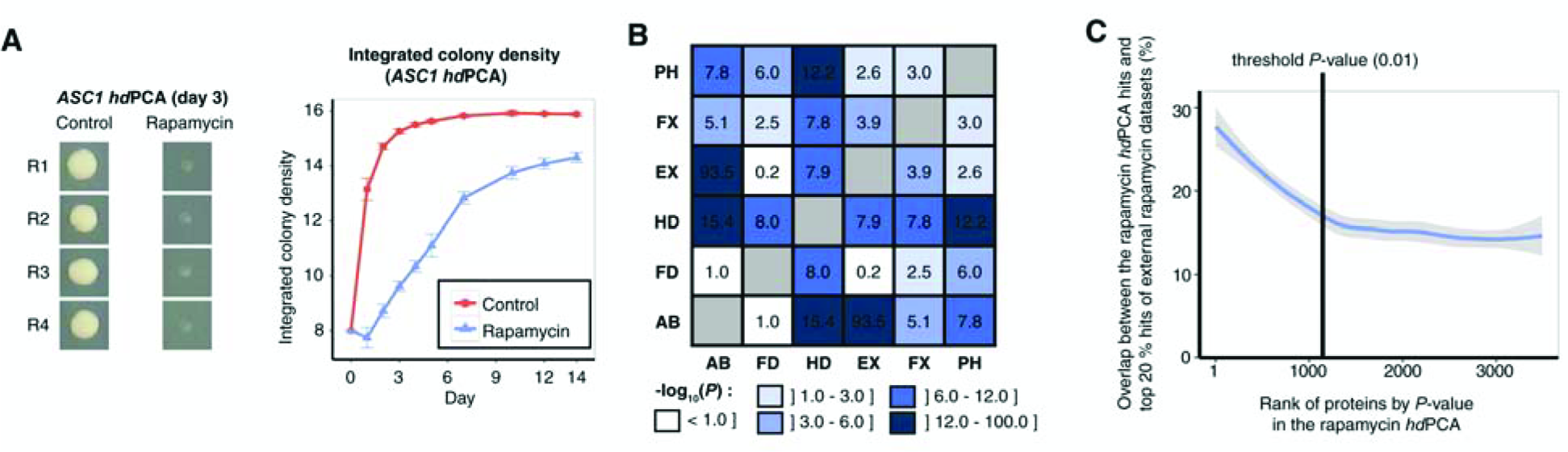
Optimization and validation of *hdPCA* with rapamycin data. **(A)** Growth of a strain containing *ASC1-DHFR-F[1,2]* and *ASC1-DHFR-F[3]* during the course of an *hd*PCA screen with rapamycin. Four replicates (R1 to R4) were tested. **(B)** Correlations among different large-scale datasets comparing control *versus* rapamycin-treated yeast cells. Data sets are compared with each other for significant overlap of genes or proteins affected by rapamycin. Values correspond to ‒log_10_(P) where *P* is the P-value of the hypergeometric test which compares significance of overlap between datasets. AB: protein abundance; FD: fitness of deletion strains; HD: *hd*PCA; EX: mRNA expression; FX: protein flux; PH: phosphoproteomics. **(C)** A LOESS curve indicating the average overlap between *hd*PCA hits, ranked by P-value, and the top 20% hits of five external rapamycin datasets (deletion screens, mRNA expression data, phosphoproteomics data, protein abundance data, and protein flux data). The enrichment found in top ranked *hd*PCA hits falls to background levels close to the threshold P-value of 0.01, which was chosen as a threshold value in further experiments.

Large-scale screens that measure individual properties of a gene, its interactions or its products are notoriously orthogonal to each other, showing little overlap in changes of individual quantities (*23*). If *hd*PCA truly captures the integrated effects of changes to a protein’s fate following a change of cellular conditions then we should expect significant correlations between *hd*PCA reporter responses and other large-scale data. We assessed how well the results of the rapamycin *hd*PCA reflect the integrated effects of the drug (Files S1-2) by comparing the data with other large-scale data on rapamycin perturbation of different gene variables (gene deletion, mRNA and protein abundance, flux, and phosphorylation levels) obtained from external sources (Fig. 2B; see Supplementary Methods for details on the datasets). We found that *hd*PCA agrees better with the other datasets than any other dataset with the exception of a strong correlation between changes in mRNA and protein abundance. This observation implies that the *hd*PCA incorporates combinations of passive (e.g., mRNA and protein abundance) and active (e.g., fitness of deletion strains, flux and phosphorylation) protein changes.

The overlap between rapamycin *hd*PCA hits (affected strains) and hits in other rapamycin large-scale studies was used to determine a criterion for a significance cutoff of differences in colony size between control‐ and drug-treated strains. At *P* > 0.01, the overlap between rapamycin *hd*PCA data and other datasets reached a plateau and hence was chosen as cutoff criterion (Fig. 2C). The significant results include 124 proteins that are essential for yeast and which are difficult to study in deletion screens.

## 3. The *hd*PCA reveals pleiotropic cellular effects by the antidiabetic drug metformin

Systematic global analyses of cellular processes that might be affected by metformin are lacking and the *hd*PCA screen provides a unique opportunity to obtain a general overview of the impact of metformin on cellular function. The *hd*PCA was carried out with metformin and identified 342 proteins with increased signal and 403 proteins with reduced signal in the presence of metformin, relative to the control condition (Files S3-4). Protein hits from the screen were more likely to modulate yeast sensitivity to metformin than proteins whose *hd*PCA signal was not significantly changed by the drug, as determined by a deletion screen (Fig. S1).

We next investigated the relationship between our observations and potential beneficial effects of metformin to T2D and to prevention of different cancers (*24*). To this end, we assessed the overlap between metformin *hd*PCA hits and human homologs containing single-nucleotide polymorphisms (SNPs) associated with T2D and with different types of cancer identified by genome-wide association studies (GWAS) that are annotated in the GWAS Catalog database (*25*). We found a significant overlap between our hits and homologous genes associated with T2D and prostate cancer (*P* = 0.01 and 0.02, respectively, hypergeometric test, Table S1). Among the eleven genes (ARL1, *CMK2, CDC55, COT1, ECM14, FYV10, HSL1, PAH1, PAB1, SFP1, SSM4*) in common between our hits and homologs associated to T2D, *SLC30A8*, the human homolog of *COT1*, was found to be associated with T2D in 12 out of 42 GWASs (Table S1 and S2). *SLC30A8* encodes a zinc transporter in the secretory vesicles of pancreatic β cells, which are implicated in insulin storage with insulin as a hexamer binding two zinc ions before secretion (*26*). A non-synonymous mutation in *SLC30A8* (R325W; rs13266634) might thus affect storage, stability or secretion of insulin in carriers of this mutation. Interpretations of the association between this mutation and the risk or prevention of T2D have been contradictory: loss of function of *SLC30A8* confers protection from T2D, whereas the decreased activity of *SLC30A8* is most commonly associated with a higher risk of developing T2D (*27-29). COT1*, the yeast homolog of *SLC30A8*, encodes a vacuolar transporter mediating zinc transport into the vacuole in a zinc replete environment and it is required for zinc and cobalt detoxification and for surviving zinc shock (*30-32). COT1* is also activated by the two transcription factors Aft1/2 on iron limitation (*33*). Cot1 exhibits an increase in signal on metformin (*P* = 1.1 x 10^−4^), explaining thus the potential benefits of metformin concordant with a higher T2D risk linked with decreased activity of *SLC30A8* in human.

One of the other overlapping genes, *FYV10*, is a negative regulator of glucose production (*34*). Notably, its human homolog, *MAEA*, is not known to be involved in this process. Its appearance in the overlap between *hd*PCA hits and T2D GWAS suggests that *MAEA* might also be involved in regulation of glucose metabolism in humans.

Recently, a growing body of evidence supports a potential therapeutic benefit of metformin in prostate cancer (*35*). The fifteen genes (*CDC8*, *DAL80*, *DID4*, *GZF3*, *MYO4*, *PRP6*, *PHB1*, *RAD51*, *RNR2*, *SCP160*, *SFP1*, *SPS1*, *SSM4*, *VMA1* and *GAT1*) in common between our hits and homologs associated to prostate cancer include two genes that are directly implicated in the cell cycle, and hence possibly in cancer: *SCP160* (increased signal on metformin, *P* = 2.4 x 10^−5^) maintains exact chromosome ploidy (*36*), and *RAD51* (increased signal on metformin, *P* = 2.1 x 10^−5^) is involved in DNA damage repair (*37*) (Table S3). In addition, among the 15 common genes between our hits and homologs associated to prostate cancer, *ATP6V1A*, the human homolog of *VMA1*, encodes the catalytic subunit of the peripheral V1 complex of vacuolar ATPase responsible for the acidification of various intracellular compartments. *ATP6V1A*, associated to the oxidative phosphorylation pathway, is overexpressed in lethal prostate cancer, while its homolog in yeast, Vma1, exhibits a decreased signal on metformin (*P* = 3.7 x 10^−4^) (*38*). In conclusion, these results follow the proposed role of metformin in the prevention of T2D (*39*) and imply a potential preventive and/or therapeutic role of metformin in prostate cancer in addition to its previous suggested preventive and/or therapeutic role in other cancers (*24, 40*).

A gene ontology (GO) enrichment analysis (Bioconductor GOstats package in R) was performed to obtain an overview of cellular processes affected by metformin (Fig. 3) (*41-43*). We used GOstats to identify biological processes that are overrepresented in proteins showing increased signal on metformin and control respectively. The enriched GO terms (*P* < 0.05) are illustrated as a network of biological processes that are organized in space according to their mutual overlap and clustered based on their relationships. The screen uncovered a wide set of GO biological processes affected by metformin and is classified into four groups (metabolism, signalling and regulation, transport and other processes) (Fig. 3, Files S5-6).

**Fig. 3.**
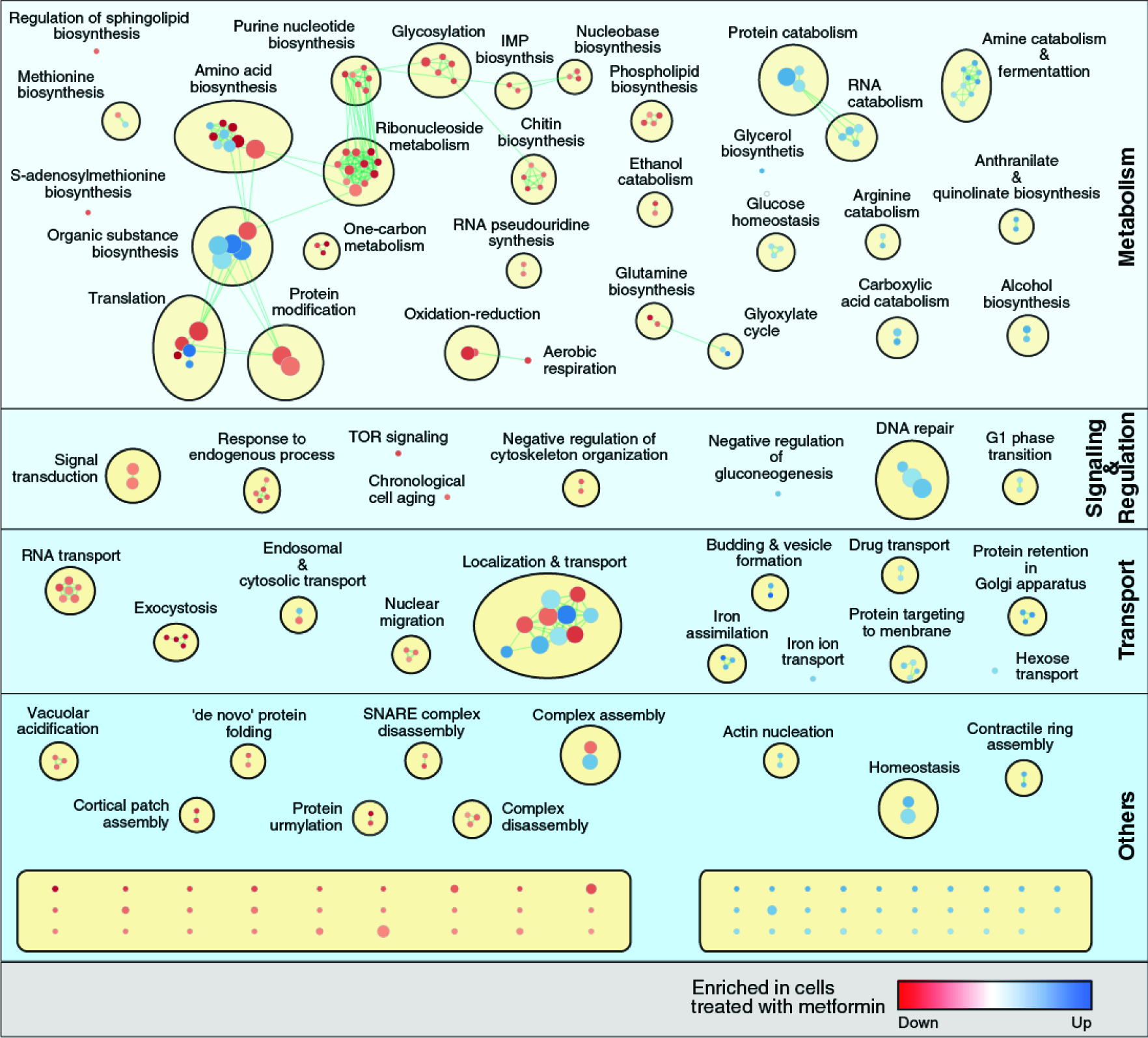
Enrichment map of GO biological processes in the metformin hdPCA. The map displays the enriched GO terms in metformin *versus* control (blue) and those that are enriched in control *versus* metformin (red). GO terms that have associated genes in common are linked with an edge. The edge width is proportional to the overlap between the linked GO terms. GO terms closer to each other in space are more functionally related than those further from each other in space and are clustered together. Ungrouped processes not mentioned in the main text are found at the bottom (unnamed; see Suppl. Tables S5 and S6 for more details). The figure is generated using the Enrichment Map (GO terms cutoff: 0.05 and Similarity measure: Jaccard Coefficient with default settings) and AutoAnnotate (default settings) plugins.

An important aspect of the antidiabetic properties of metformin originates from its ability to interfere with energy metabolism. Metformin stimulates glucose uptake, reduces gluconeogenesis and cellular respiratory capacity, and increases concentrations of glycerol and lactic acid (reviewed in (*44)*). In accordance with these observations in higher eukaryotes, the *hd*PCA in yeast shows an increased signal for the major glucose transporters, for proteins involved in breakdown of gluconeogenic enzymes and for enzymes involved in glycerol and ethanol production (Fig. 3; Fig. 4A). In contrast, the *hd*PCA signal of members of the citric acid cycle and oxidative phosphorylation is reduced. This cross-species conservation of the effect of metformin on central carbon metabolism was previously reported in a study that showed increased glucose uptake and glycerol production in yeast despite its lack of a structurally conserved mitochondrial complex I, the most cited potential target of metformin (*45*). Indeed, sensitivity to metformin follows the degree to which yeast depends on respiration. Higher levels of the fermentable sugar glucose reduce the effect of metformin on yeast growth while carbon source conditions that rely partially (galactose and low glucose) or completely (glycerol) on respiration increase sensitivity to metformin (Fig. 4B). Furthermore, the proton gradient across the mitochondrial membrane is reversed after prolonged metformin exposure (Fig. 4C), indicative of an interference with ATP production based on the electron transport chain, the final phase of respiration. Even though the mitochondria are considered to be the main target site of metformin, the effect of this drug on cellular processes extends far beyond this. For example, proteins involved in one-carbon and nucleotide (dNTP) metabolism are negatively affected by metformin according to the *hd*PCA data (Fig. 4D), which is in line with previous observations (*46, 47*). Nucleotide synthesis requires methyl groups from folate intermediates produced in one-carbon metabolic processes. We confirmed by mass spectrometric analysis that dNTP levels are reduced in yeast upon metformin treatment (Fig. 4E). Overall, catabolic processes tend to be positively affected by metformin while *hd*PCA reporters involved in anabolic processes are reduced in the presence of the drug (Fig. 3).

**Fig. 4.**
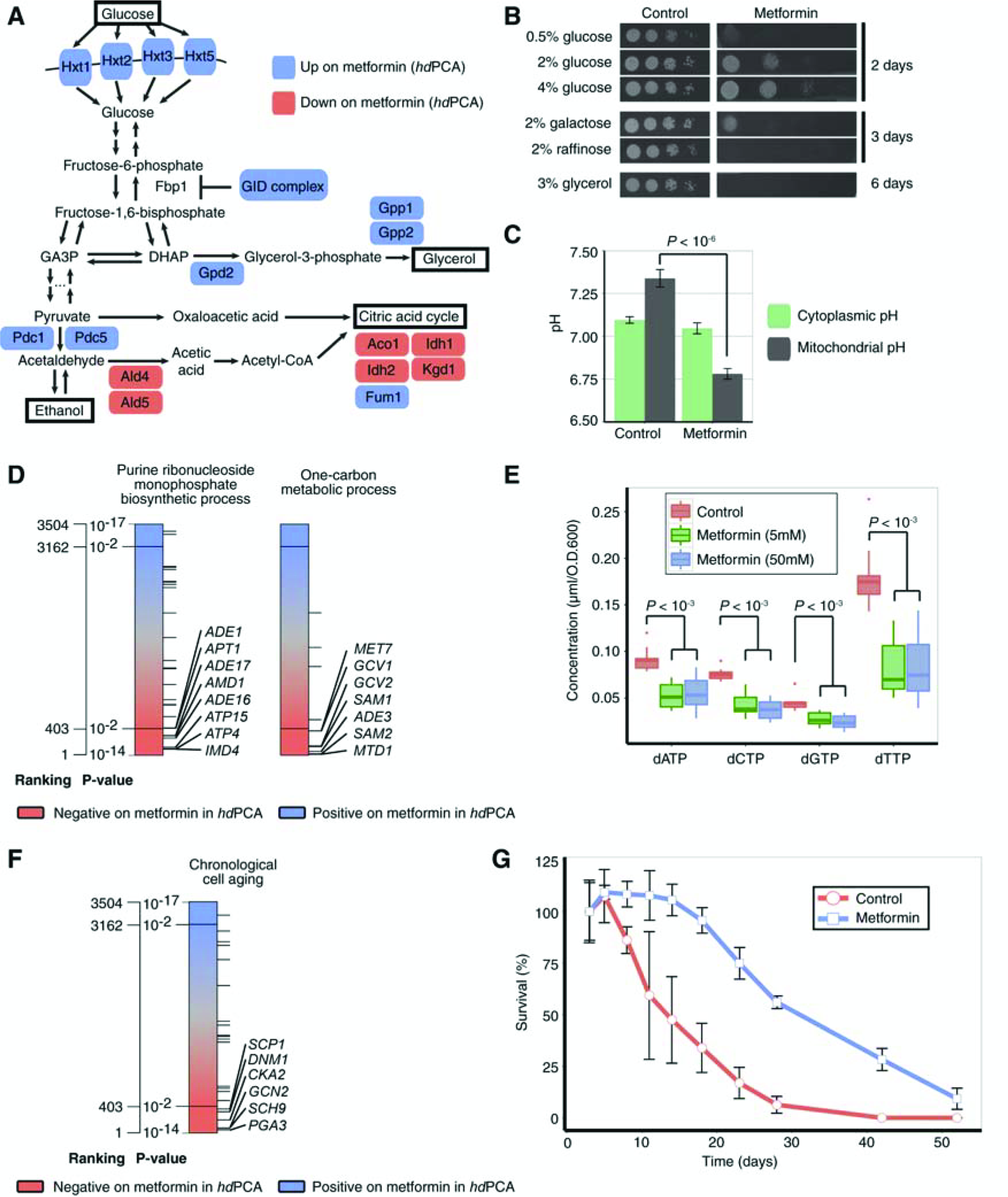
Metformin influences a range of cellular processes. **(A)** Metformin increases (blue) and decreases (pink) the *hd*PCA signal of proteins involved in glucose uptake and catabolism. **(B)** Sensitivity to metformin changes with carbon source. Pictures were taken on different days to account for the basal effect of carbon source on growth. **(C)** The mitochondrial membrane proton gradient reverses after prolonged metformin treatment (7h). The pH was determined using a pH-dependent fluorescent protein (pHluorin) with or without mitochondrial localization sequence. **(D)** The *hd*PCA signals for proteins involved in purine ribonucleoside biosynthesis and one-carbon metabolism are reduced in the presence of metformin. Each horizontal line represents one protein member of the biological process. **(E)** Metformin at 5 mM and 50 mM reduces the concentration of dNTPs. Concentrations were determined by mass spectrometric analysis of the individual dNTPs (12 replicates for each condition). **(F-G)** Metformin prolongs chronological lifespan. Three cultures were incubated in the presence of metformin and three in its absence. Cell survival was tracked over time by counting the colony-forming units at regular time points. The error bars correspond to the standard deviation of three replicates of one qualitatively representative experiment. GID: glucose-induced degradation; GA3P: glyceraldehyde-3-phosphate; DHAP: dihydroxyacetone phosphate.

On the level of cell regulation, three important results appear from the *hd*PCA. First, proteins that stimulate chronological aging show a reduced *hd*PCA signal on metformin (Fig. 4F), consistent with observations of increased yeast longevity after metformin treatment (*45*). We found that metformin increases the median survival time by 16 days, a 3.2-fold increase compared to the control condition (Fig. 4G). The second form of regulation that appears in the *hd*PCA data is the TOR pathway exhibiting reduced signals for several of its associated members in the presence of metformin (Fig. S2A). In breast cancer cells, metformin inhibits the TOR pathway and reduces translation rates (*48*). Moreover, the kinase Sch9, which is tightly linked to TOR signaling, is the most significant hit for a major regulatory protein, with a decreased *hd*PCA signal on metformin (*P* = 2.04 x 10^−6^). Interestingly, its human homolog, S6 kinase, is inhibited by metformin (*48*). Finally, the *hd*PCA data suggest that metformin induces a DNA repair response in yeast. An understanding at the molecular level of the relationship between metformin and DNA repair is of particular interest for alternative applications of metformin administration in cancer prevention and cancer chemotherapy. Our data supports the involvement of the *SWI/SNF* and *INO80* chromatin remodeling complexes in metformin-mediated DNA repair mechanisms (Fig. S2B).

## 4. Metformin provokes a cellular state akin to iron deficiency

Interestingly, a common theme appears upon global analysis of the affected cellular processes in yeast upon metformin treatment. Reduction in respiratory capacity, increased glucose uptake, increased glycerol production, inhibition of the TOR pathway, increased life span and activation of DNA repair are all found under conditions of iron limitation in yeast and/or other organisms (Fig. 5A) (*49-54*). Furthermore, other cellular functions found in the *hd*PCA data such as RNA transport, vesicle-mediated transport, the glyoxylate cycle and folate metabolism have been linked in the past with reduced iron availability(55-58). In addition, iron-binding proteins show a reduced *hd*PCA signal on metformin (Fig. 5B), whereas proteins important for resistance to iron limitation tend to have a positive signal in the *hd*PCA (1.7-fold enrichment, *P* = 2.9 x 10^−4^) (*56*).

**Fig. 5.**
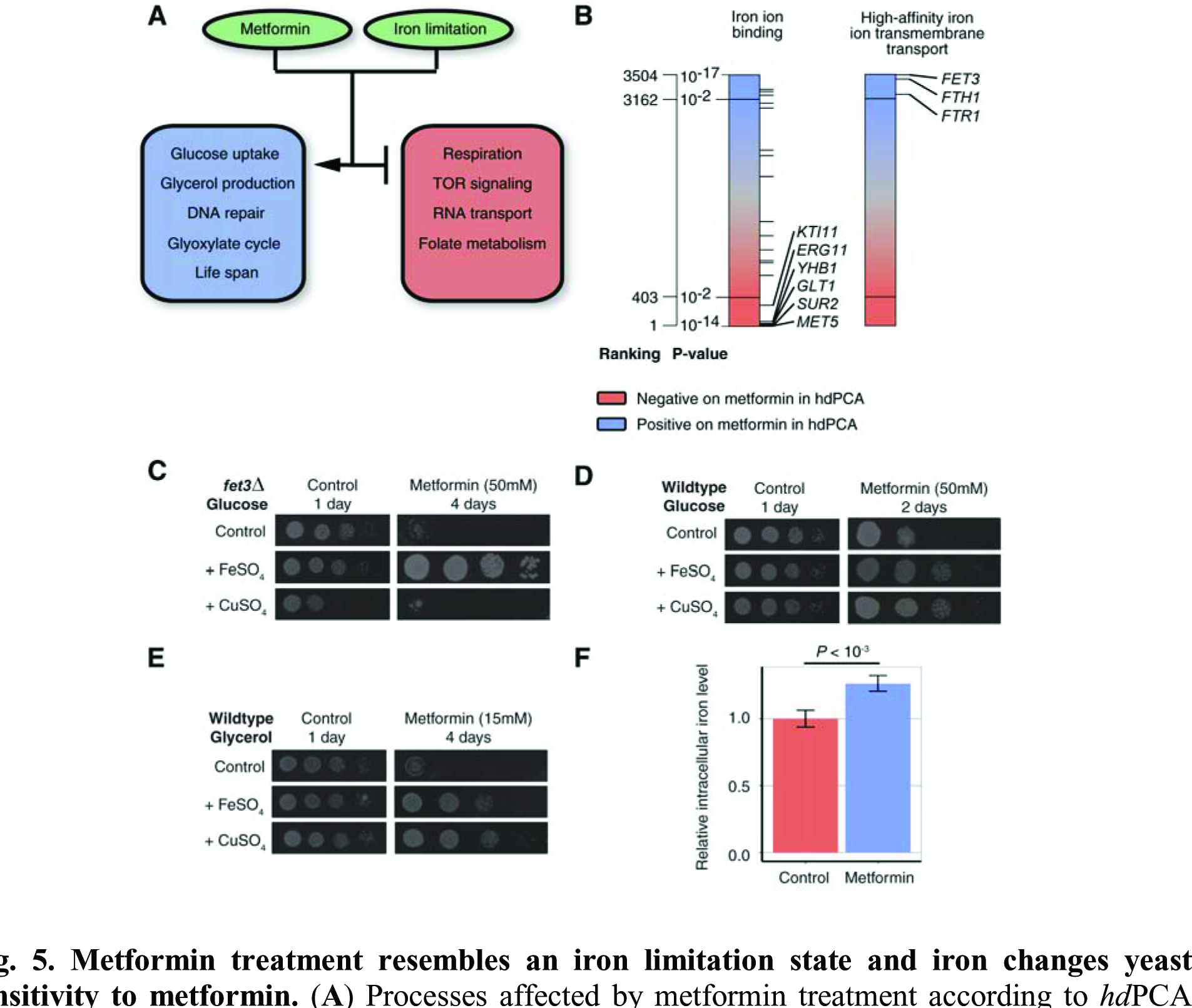
Metformin treatment resembles an iron limitation state and iron changes yeast sensitivity to metformin. **(A)** Processes affected by metformin treatment according to *hd*PCA data are also modulated by iron limitation. **(B)** Metformin *hd*PCA data on proteins involved in high-affinity iron ion uptake and proteins binding iron. **(C)** Growth of *fet3Δ* in the presence of metformin and CuSO_4_ or FeSO_4_. **(D)** Growth of wild-type yeast in the presence of metformin and CuSO_4_ or FeSO_4_. **(E)** Growth of wild-type yeast in respiratory conditions with metformin and CuSO_4_ or FeSO_4_. **(F)** Intracellular iron levels of cells in stationary phase in control and metformin conditions.

An increase in *hd*PCA signal for the high-affinity iron uptake system provides a more direct link between the *hd*PCA data and iron deficiency (Fig. 5B). Yeast becomes more sensitive to metformin upon deletion of the ferro-O2-oxidoreductase *FET3*, which is directly involved in copper-dependent high-affinity iron uptake (Fig. 5C). Iron supplementation overcomes the growth deficiency of *fet3Δ* in the presence of metformin. Furthermore, addition of iron (FeSO4) or copper (CuSO_4_) to growth medium makes wild-type yeast less sensitive to the drug (Fig. 5D). The antagonistic effect of copper on metformin-mediated growth inhibition requires Fet3, which suggests that copper mainly improves yeast growth by stimulation of copper-dependent iron uptake. Under respiratory growth conditions, which require more iron than fermentable conditions, the supplementation of iron (or copper) counters the effects of metformin on growth more profoundly (Fig. 5E). These results suggest that metformin either interferes with iron uptake or with intracellular iron homeostasis. We found that metformin does not reduce the intracellular iron content during exponential growth (Fig. S3) and even increases intracellular iron levels in stationary phase cultures (Fig. 5F). This result indicates that metformin generates a condition perceived by the cell as iron deficient even though there is no shortage of iron inside the cell.

## 5. Conclusions

Our observations suggest that metformin somehow acts upon sensors of iron levels (perhaps Fet3) to signal an iron deficient state to the cells. Iron supplementation partially compensates for this perturbation. These results may bear directly on observations in humans. Iron excess caused by hereditary factors, nutrition or medical intervention is a diabetes risk factor (reviewed in (*59*)). A low-iron diet or iron chelation therapy improves the glycemia of diabetes-prone, leptin-deficient mice (*60*). Metformin treatment leads to a trend of reduction in serum iron levels (*61*). Our results warrant a closer investigation of the effects of metformin on iron homeostasis, accessibility and distribution at the possible sites of metformin activity, including the gut and the liver.

With hdPCA, we were able to uncover the effect of metformin on a broad set of cellular processes and we found cross-species conservation of core mechanisms of metformin treatment in yeast, such as its effect on central carbon metabolism, nucleotide metabolism and longevity. This opens up the possibility for future research on metformin with this model organism. Furthermore, the overlap between the *hd*PCA data and GWA studies related to diabetes and prostate cancer narrows down the human genes of interest in establishing important predictors of diabetes risk and genes involved in manifestation of prostate cancer.

Finally, *hd*PCA provides a simple strategy to globally profile cellular responses to external perturbations that can be applied widely to study mechanisms of action of bioactive molecules, their effects on intended target or unsuspected off-target effects. Identifying off-target effects are particularly important for determining both potential liabilities and added benefits of molecules (*62*). Applications of *hd*PCA include the general study of the impact of environmental stresses, revisions of mechanisms of action of commercialized drugs, comparison of off-target effects within a drug family and elucidation of the harmful effects of toxins.

## Acknowledgements

We would like to thank Dr. Gertien Smits for pHluorin plasmids. Supported by the Canadian Institutes of Health Research (MOP-GMX-152556 and MOP-GMX-231013) (S. W. M.), Natural Sciences and Engineering Research Council (05707-2015) (S. W. M.), Human Frontier Science Program (RGP0050/2013) (S. W. M.), the Francis Crick Institute from Cancer Research UK (FC001134) (M. R.), the UK Medical Research Council (FC001134) (M. R.), the Wellcome Trust (FC001134 and RG 093735/Z/10/Z) (M. R.) and the ERC (Starting grant 260809) (M. R.).

## Supplementary Materials

Materials and Methods Table S1 - S3 Fig. S1 - S2

Additional Data Tables S1 - S10

